# The HP1γ epigenetic silencer dampens IFN-γ response at the gut epithelial barrier

**DOI:** 10.1101/2022.12.27.522038

**Authors:** Yao Xiang, Jorge Mata-Garrido, Christophe desterke, Eric Batsché, Ahmed Hamaï, Youssouf Sereme, David Skurnik, Abdelali Jalil, Jean-Christophe Beche, Eliane Piaggio, Laurence Arbibe, Yunhua Chang

**Author notes:** Co-authorship (Laurence Arbibe, Yunhua Chang).

## Abstract

Interferon gamma (IFN-γ) plays central roles in the pathophysiology of inflammatory bowel disease (IBD), both activating inflammatory responses and immunosuppressive functions. However, the epigenetic mechanisms controlling the expression of IFN-γ responsive genes at the gut epithelial barrier are not well understood. In this study, we identified the epigenetic regulator HP1γ as a transcriptional repressor of the IFN-γ-responsive genes STAT1 (signal transducer and activator of transcription 1) and PD-L1 (Programmed Cell Death Ligand 1). Accordingly, HP1γ gene inactivation in the mouse gut epithelium resulted in an immunopathology with a long-lasting up-regulation of STAT1 and PD-L1. *C*olon organoids models and i*n vitro* cell lines showed that HP1γ deficiency primed STAT1 and PD-L1 expressions, ultimately sensitizing epithelial cells to IFN-γ stimulation. Chromatin immunoprecipitation experiments suggest that HP1 promoter tethering is involved in the silencing of gene expression. Overall, these results identify HP1γ as an epigenetic silencing pathway controlling the IFN-γ response at the epithelial barrier.

## Introduction

Inflammatory bowel disease (IBD), including ulcerative colitis (UC) and Crohn’s disease (CD), are chronic inflammatory gut disorders characterized by a deregulated mucosal immune response. Cytokines released by different pro-inflammatory T cell types are central mediators of the lesions in the inflamed intestinal mucosa of IBD patients. While initial studies have identified Th1/Th17 lineage cells enriched in CD patients (Weaver et al., 2013), recent studies have extended these observations to UC (Lamb et al., 2017) (Iacomino et al., 2020). Notably, high level of IFN-γ was found in the colonic epithelial surface of UC patients (Iacomino et al., 2020). In response to IFN-γ, the activator of transcription (STAT) become activated after phosphorylated by Janus kinases (JAK). The activated protein migrates into the nucleus and binds to specific promoter elements to regulate gene expression. Many immunoregulatory genes contain multiple STAT1 binding sites, including *STAT1* itself and Interferon regulatory factor 1(*IRF1*). The later in turn can promote transcription of *CD274* (Programmed death-ligand 1, PD-L1), the main ligand for the T cell surface receptor PD-1, a negative co-stimulatory molecule inhibiting T cell function (Han et al., 2020). Likewise, an up-regulation of the mRNA encoding for PD-L1 has been evidenced in the IBD mucosa, suggesting a possible protective function by dampening aberrant T cell activation in the course of the disease (Rajabian et al., 2019).

Increased expression of STAT1 has been documented in IBD, particularly in UC as well as in mice model of DSS-induced colitis (Schreiber et al., 2002). Yet, the regulatory mechanisms controlling the expression of STAT1 and its downstream target genes in the intestinal epithelium remains poorly understood. In an earlier study, we reported that the expression of the chromatin and alternative splicing regulator HP1γ was reduced in UC patients (Mata-Garrido et al., 2022). Importantly, HP1γ depletion in the mouse gut epithelium triggers IBD-like traits, including inflammation and dysbiosis. We now showed that HP1γ controls *in vivo* the IFN-γ signaling cascade in the colon epithelium by directly repressing STAT1/PDL1 expressions. Likewise, HP1γ deficiency sensitizes epithelial cells to IFN-γ signaling, suggesting a potential of HP1γ in regulating appropriate immune response at the colon mucosa.

## Results

### *Cbx3* KO mice exhibit long-lasting goblet cell alteration together with immunopathology

As Goblet cell alterations predispose to colitis (Nyström et al., 2021), we firstly documented goblet cell/mucin production by Periodic Acid Schiff staining in Villin-creERT2:*Cbx3*^-/-^ mice 7 days and 12 months post-tamoxifen inductions (**Figures 1A-C**). The staining revealed a potent alteration in the detection of the signal that was maintained over-time (**Figure 1B**–**1C**). Crypt lengths, a proxy of inflammation-induced regeneration, were chronically increased at both proximal and distal colons (**Figure 1D**). Importantly, CD4^+^ and CD8^+^ T cells were significantly increased in the proximal and distal colons of the *Cbx3* KO mice at day 7 post-tamoxifen induction (**Figure 1E**–**1F**). Remarkably, infiltration of CD4^+^ T cells in the proximal colon was long-lasting, as shown by the increased detection in the mucosa of the 12 months aged *Cbx3* KO mice (**Figure 1F**). Likewise, the size of lymphatic nodules appeared larger in *Cbx3* KO mice as compared to WT mice (**Figures 1G-H**). Overall, these data indicate that HP1**γ** deficiency triggers long-lasting colon mucosal alterations, dominated by goblet cell alterations and a chronic activation of the immune response.

**Figure1:**
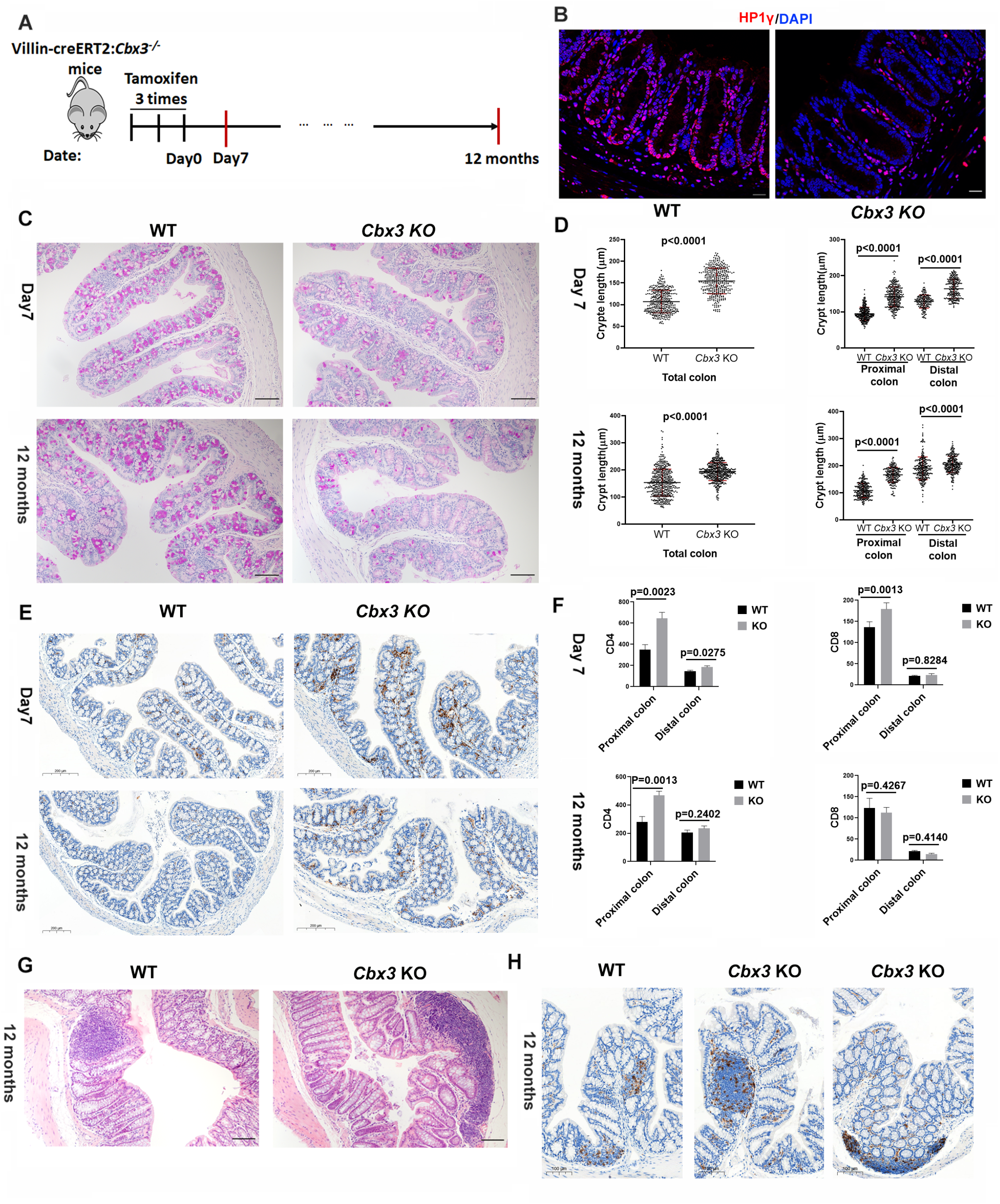
**(A)** Scheme illustrating the time-course of sample collections. Samples were collected upon *Cbx3* knock-down 7-day and12 months post-tamoxifen gavage **(B)** Representative immunofluorescence in colon tissue sections from WT and *Cbx3*^-/-^ mice with anti-HP1γ antibody (red) and DAPI (blue).Scale bar: 20μm. **(C)** Representative sections stained with PAS (Periodic Acid Schiff) in WT and *Cbx3* KO colon tissues at the indicated ages. **(D)** Colon crypt lengths measured by Image J, 4 mice in each group, two-sided Student’s t-test) (**E)** Representative immunochemistry with anti-CD4 antibody in WT and *Cbx3* KO colon tissues. Hematoxylin (blue) counterstaining was used for nuclear coloration. **(F)** CD4^+^ or CD8^+^ cells counting in WT and *Cbx3* KO colon tissues 7 days or 12 months post-tamoxifen gavage. Y axis represents CD4^+^ or CD8^+^ cell number counted in each 4μm thickness colon section from proximal or distal colon (4 mice in each group, two-sided Student’s t-test). **(G and H)** Representative HE (Hematoxylin and eosin) staining revealing an increase size in lymphoid nodule in 12 months *Cbx3* KO mice colon. Scale bar: 20μm.

### HP1γ targets STAT1 and PD-L1 genes in the mice colon epithelium

We next sought to identify immune genes directly targeted by HP1. To that end, we crossed the RNA-seq data obtained from purified *Cbx3* KO mice colon epithelium ((Mata-Garrido et al., 2022), GSE192800) with a publicly available HP1γ Chromatin immunoprecipitation-sequencing (ChIP-seq) performed in the HCT116, a colorectal cancer cell line ((Smallwood et al., 2012), GSE28115). This strategy led us to identify 68 genes as potential HP1γ target genes (**Figure 2A**). Interestingly, among the group of up-regulated genes by *Cbx3* inactivation, the STRING network analysis revealed a cluster composed by 10 HP1γ direct targets implicated in the immune responses. Among them, the string node identified signal transducer and activator of transcription STAT1 (signal transducer and activator of transcription) and CD274 (Programmed cell death 1 ligand 1, PD-L1), two components of the IFN-γ signaling pathway (Figure **2B**). On these genes, ChIP-seq HP1γ evidenced peaks of HP1γ associations at promoter regions (**Figure 2C-D**) and RNA seq revealed that transcription was significantly up-regulated upon *Cbx3* inactivation (**Figure 2E-F**). In parallel, GSEA analysis revealed significant enrichments of genes involved in the inflammatory/ IFN-γ responses in the *Cbx3* KO colon epithelium (**Figure 2G**). Likewise, MHCI/II (Major histocompatibility complex class I/II) and chemokines gene expressions were significantly increased (**Figure 2 H-J).**

**Figure 2:**
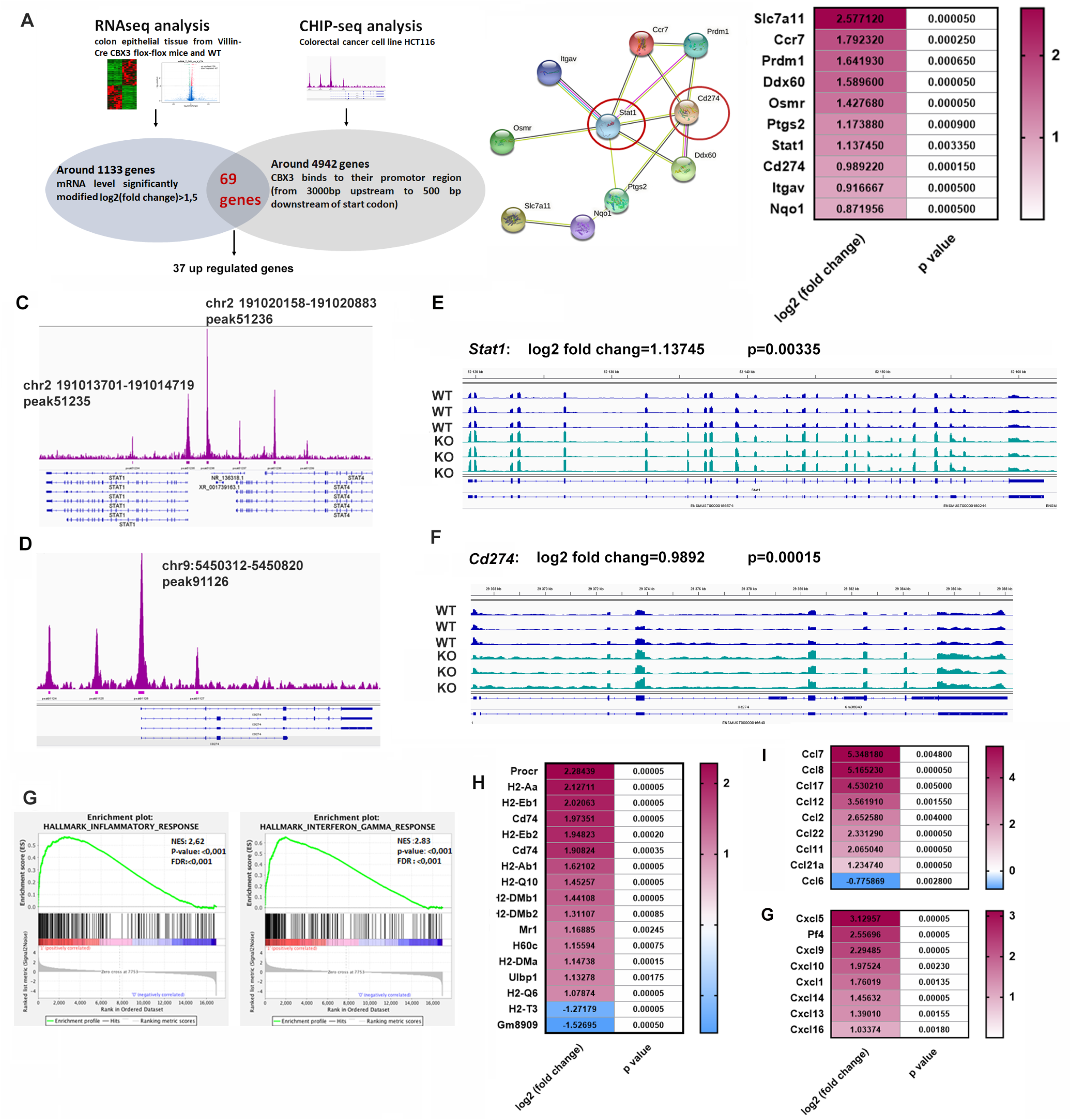
**(A)** Workflow for identifying HP1 target genes: RNA sequencing revealed 1133 genes whose expression level extends more than 1.5 fold (increase or decrease) in *Cbx3*^-^KO colon epithelium, as compared to WT. The ChIP-seq identified 4942 genes with proximal promoter region enrichment for HP1γ (from 3000bp upstream to 500bp downstream of the Transcription Starting Site (TSS). Crossing the 2 dataset led to the identification of 68 genes (37 upregulated and 31 down-regulated genes) as potential HP1γ target genes.**(B)** String analysis revealed a cluster composed by HP1γ direct targets implicated in the immune responses. Among them, the string node identified signal transducer and activator of transcription STAT1 and CD274 **(C and D)** ChIP-seq analysis identified two HP1γ binding sites at the *STAT1* promoter (**C**) and one at the *CD274* promoter (**D**). **(E and F)** Integrative Genomics Viewer (IGV) was applied to visualize increased *Stat1* and *Cd274* transcription levels in 3 *Cbx3*KO mice compared to 3 WT mice. (**G)** GSEA analysis based on transcriptome data revealed a significant enrichment of inflammatory and IFN-γ responses in *Cbx3*KO colon epithelium. **(H-J)** Heat map showing centered on IFN-γ responsive genes. MHCI/II (Major histocompatibility complex class I/II) and chemokines gene expressions were significantly increased in *Cbx3*KO colon epithelium, as compared with WT.

To validate bioinformatic analysis, the expressions of STAT1 and PD-L1 were determined in *Cbx3* KO mice 7 days and 12 months post-tamoxifen induction. Increased STAT1 and PD-L1 protein levels were detected in *Cbx3*^-/-^ colon epithelium at both time points (**Figure 3A-B**). Likewise, immunofluorescence studies revealed an intense STAT1 nuclear labeling in the colon epithelium of the *Cbx3* KO mice (**Figure 3D–3E**). Importantly, Q-PCR data showed that HP1 durably represses mRNA expressions for *Stat1* and *Pd-l1* in the colon epithelium (**Figure 3C).** Overall, these results spotlight HP1γ as a repressor for STAT1 and PD-L1 expressions in the colon epithelium.

**Figure 3:**
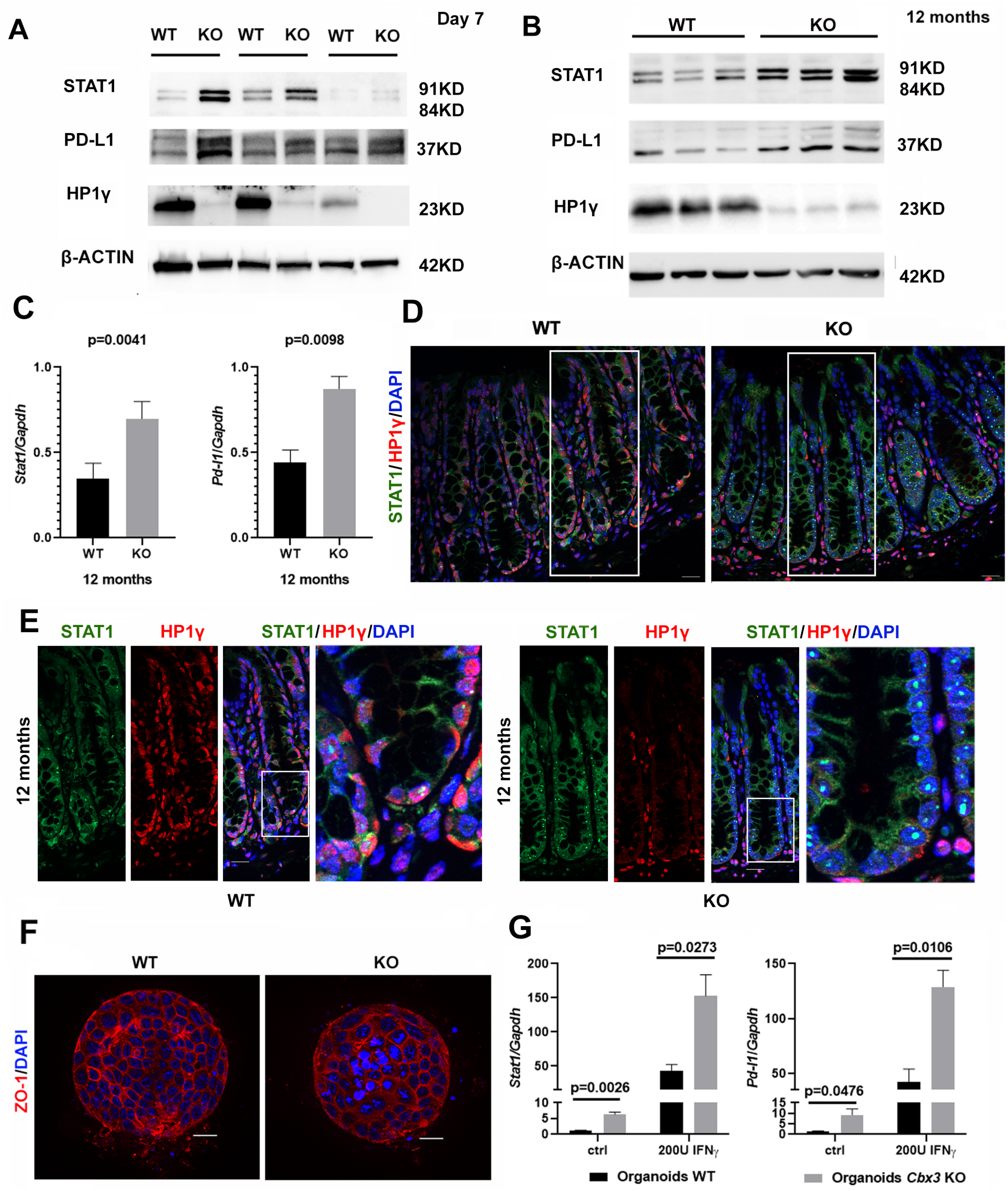
(**A-B**) Immunoblots STAT1 and PD-L1 in colon crypts epithelium 7 days post-tamoxifen induction (n=3 mice in each group)(**C)** Western blot and RT-qPCR showing increased STAT1 and PD-L1 protein and mRNA levels in colon 12 months post-tamoxifen induction (n=3 mice in each group). (**D-E)** Representative immunofluorescence in colon tissue sections stained with anti-HP1γ antibody (red), anti-STAT1 (green) and DAPI (blue) in WT and *Cbx3*KO mice 12 months post-tamoxifen induction. The Figure 3E represents the insert from Figure 3D. **(F)** Immunofluorescence with anti-ZO-1 (red) and DAPI (blue) in organoids derived from WT and *Cbx3*KO colons marks the cell-cell tight junction **(G)** Organoids were stimulated or not by 200UI during 24h hours. mRNA expression for *Stat1* and *Pd-l1* were detected Q-PCR (n=3 mice in each group, two-sided t test).

### HP1γ deficiency primes STAT1 and PD-L1 expressions thus reinvigorating response to IFN-γ signaling

We previously showed that the microbiota of *Cbx3* KO mice was dysbiotic, with the overrepresentation of colitogenic bacterial species that may trigger the expression of interferon responsive genes in *Cbx3* KO mice (Mata-Garrido et al., 2022). To circumvent this possibility, we generated colon organoids derived from WT or *Cbx3* KO mice and stimulated them *in vitro* with IFN-γ (**Figure 3F**). Q-PCR data showed a significant increase in the basal expression of *Stat1* and *Pd-l1* gene expression in *Cbx3* KO organoids, further enhanced by addition of IFN-γ (**Figure 3G**). To rule out the possibility that an imprinting of the dysbiotic microbiota might contribute to the phenotype observed in organoids, we generated Crispr/Cas9 *CBX3* KO cells in the SW480 and HT29 human colorectal cancer (CRC) cell lines. Editing efficiency of the Crispr/Cas9 was checked by Q-PCR and Western blot analyses, with compensatory mechanisms revealing by an increase mRNA expressions for HP1α and HP1β in SW480 but not HT29 Crispr/Cas9 *CBX3* KO cells (**Figures 4A-B** and **Figure S1**). Reminiscent of the phenotype observed in the organoid model, HP1 deficiency was sufficient for increasing expression of STAT1 at both mRNA and protein levels in the 2 cell lines (**Figures 4C-D).** Remarkably, while stimulation by IFN-γ barely affected STAT1 expression in control cells, IFN-γ treatment dramatically increased STAT1 mRNA and protein expression levels in both *CBX*3 KO cell lines (**Figure 4C–4D, left and middle panels**). Immunofluorescence analysis using a phospho-STAT1 antibody further identified an increase detection of active STAT1 specifically in *CBX*3 KO cell lines (**Figure 4C–4D, right panels)**. PD-L1 expression followed a similar trend, with an increase expression level in response to IFN-γ treatment specifically observed upon *CBX3* inactivation, as shown by Q-PCR and flow cytometry analyses (**Figures 4E-F**). Overall, these data showed that HP1γ deficiency is sufficient for priming STAT1 and PD-L1 expressions, ultimately sensitizing normally unresponsive CRC cell lines to IFN-γ.

**Figure 4:**
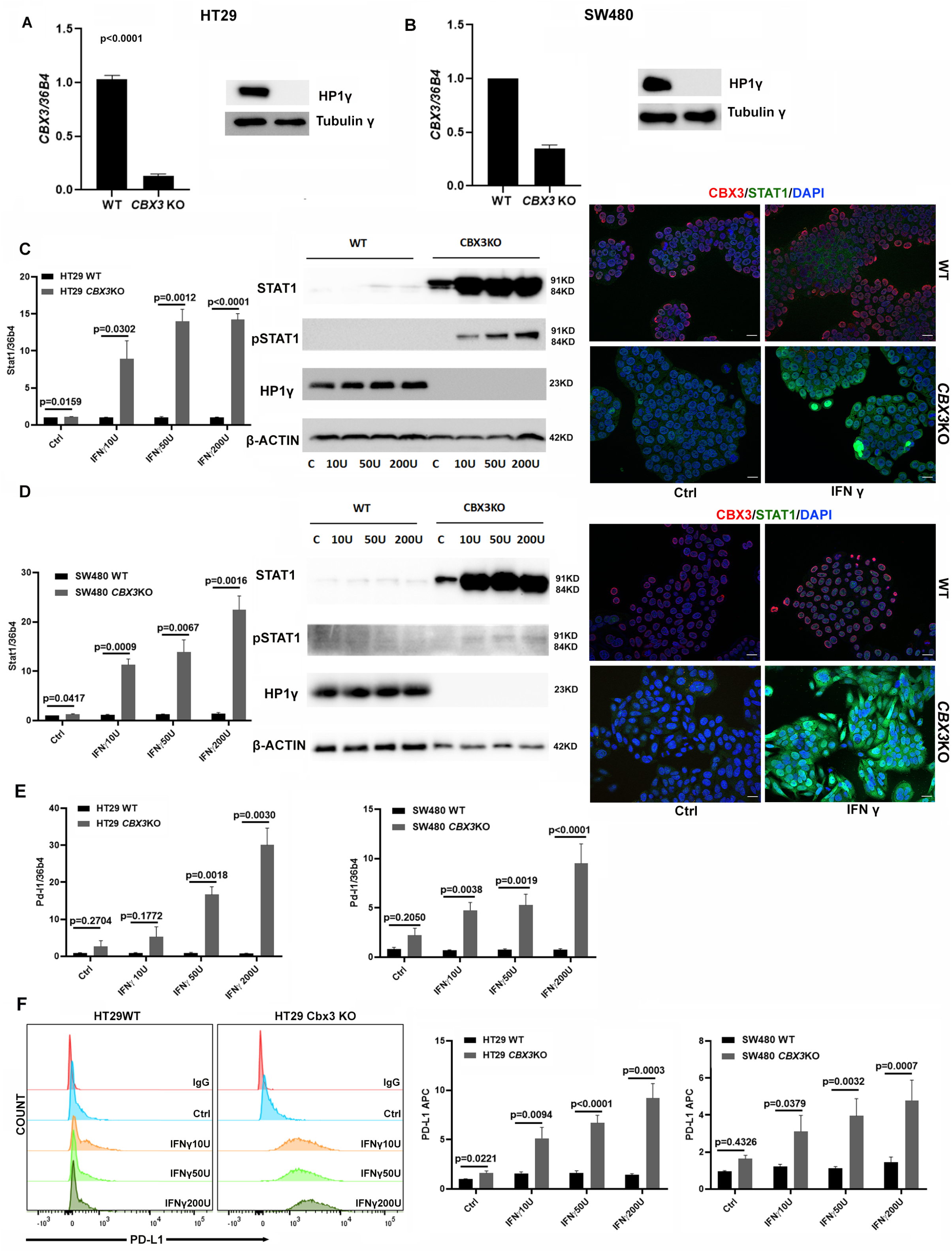
(**A**-**B**) mRNA expression for *CBX3* and immunoblot HP1γ in WR and Crispr/Cas9 mediated *CBX3* deletion in two CRC cell lines (3A: HT29 and 3B: SW480). (**C**-**D**) QPCR data (left panel), Western-blot (middle panel) and IF (right panel) revealing that IFNγ stimulation dramatically increased STAT1 expression in Crispr/Cas9 *CBX3*KO cells compared to WT control (4C: HT29 and 4D: SW480), two-sided t test for statistical analysis, n=3 independent experiments. The immunofluorescence was performed with anti-HP1γ antibody (red), anti-STAT1 (green) and DAPI (blue) for WT and KO cells without (Ctrl) or with IFNγ stimulation (200U). Scale bar: 20μM **(E)** mRNA expression for PD-L1 in HT29 and SW480 *CBX3*KO, cells compared to WT control, n=3 independent experiments, two-sided Student’s t-test (**F)** Flow cytometry analysis detecting cell surface PD-L1 expression: PD-L1 expression was significantly increased in HT29 and SW480 *CBX3*KO cells, as compared to WT upon different concentrations of IFN-γ stimulation. Left panel: Representative histogram of HT29 WT and *CBX3KO* cells is shown. Middle and right: the histograms showed the mean values of PD-L1 APC fluorescence intensity under different experimental conditions. n=3 independent experiments, two-sided Student’s t-test

### HP1γ targeting at the *STAT1* and *PD-L1* promoters is disrupted by IFN-γ stimulation

We next investigated whether HP1γ associated to the chromatin of its target genes STAT1 and PD-L1. Based on the ChIP-Seq HP1γ data, peaks of HP1γ associations were detected at promoter regions of *STAT1* and *PD-L1* genes. Accordingly, UCSC Genome Browser revealed that the potential promoter binding sites for HP1γ were located at H3K4Met, H3K27Ac and CpG islands enriched regions (**Figures 5A** and **5D** for *STAT1* and *PD-L1*, respectively).

**Figure 5:**
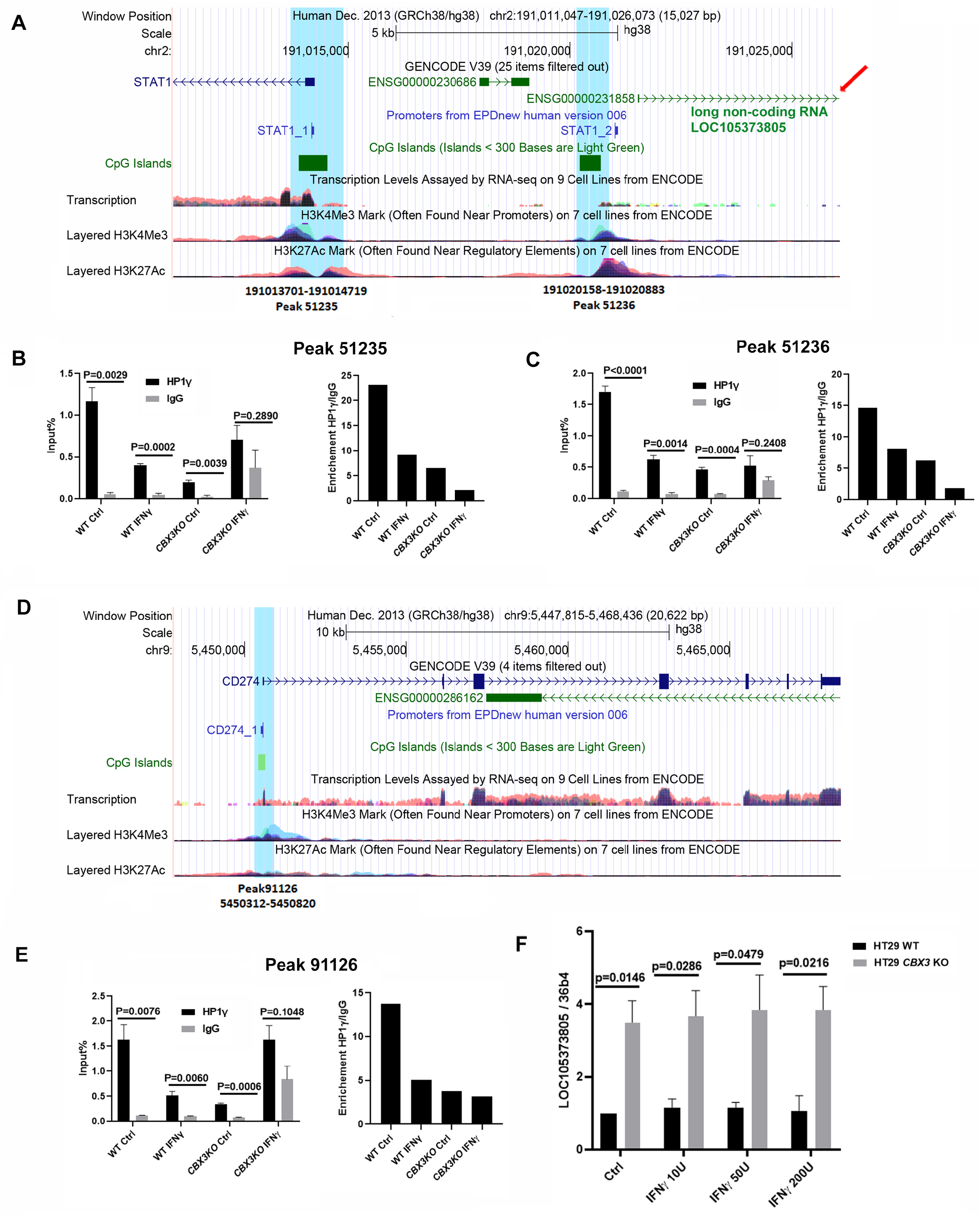
**(A)** UCSC Genome Browser centered on *STAT1:* The blue labeling corresponds to the ChIP-seq Peaks 51235 and 51236. These HP1γ binding sites are located at the promoter region of *STAT1* enriched by H3K4Me3, H3K27Ac, CpG islands. The red arrow designed the non-coding RNA LOC105373805, upstream of the *STAT1* promoter and in opposite orientation.**(B and C)** ChIP experiments were carried out with HP1γ or IgG antibodies. Input (left panel) or enrichment (right panel) relatively to the signal obtained for ChIP using non-immune IgG at Peak 51235 and 51236 regions of *STAT1* gene. Experiments were performed in WT and *CBX3* KO HT29 cell lines, stimulated or not by IFNγ. n=3 independent experiments, two-sided Student’s t-test. **(D)** UCSC Genome Browser centered on CD274 (or *PD-L1):* ChIP-seq Peak 91126 for *PD-L1* gene (CD274) is labelled in blue, matching with the promoter regions of *PD-L1*, indicated by an enrichment of H3K4Me3, H3K27Ac and CpG islands. (**E)** ChIP experiments were carried out with HP1γ or IgG antibodies. Input (left panel) or enrichment (right panel) relatively to the signal obtained for ChIP using non-immune IgG at Peak 91126 region of *CD274* gene. Experiments were performed in WT and *CBX3* KO HT29 cell lines, stimulated or not by IFNγ. n=3 independent experiments, two-sided Student’s t-test. **(F)** QPCR detecting the transcript level of LOC105373805 in WT and *CBX3* KO HT29 stimulated or not by IFNγ. n=3 independent experiments, two-sided Student’s t-test.

ChIP assays for HP1γ were performed in the WT and *CBX3* KO HT29 cell lines, with and without IFN-γ stimulation. Detection of HP1 chromatin association was performed using primers specific to the identified binding sites at peaks 51235 and 51236 for *STAT1* and peak 91126 for *PD-L1* (**Figure 5A** and **Table S2**). ChIP assays HP1γ in unstimulated WT cells revealed a potent association of HP1 to promoter regions for both genes (**Figures 5B-C and E**, left panels). Interestingly, IFN-γ stimulation led to a marked decrease chromatin association in WT cells (**Figures 5B-C and E**, left panels). By contrast, there was a basal signal insensitive to IFN-γ stimulation in the *CBX3* KO cell line (**Figures 5B-C and E**). Consistently, the average fold enrichment (HP1γ antibody/ IgG control) was highest in unstimulated WT cells, as compared to IFN-γ-treated cells (**Figure 5B-C and E**, right panels). We also noted an uncharacterized long non-coding RNA (LOC105373805), whose location was upstream of the *STAT1* promoter in opposite orientation, most likely resulting from a bidirectional STAT1 promoter activity (**Figure 5A**, red arrow). Q-PCR detection of LOC105373805 transcript level indicated that *Cbx3* inactivation significantly increased RNA expression, with no further enhancement induced by **I**FN-γ stimulation (**Figure 5F**). Thus, HP1 exerts a basal repression at the *STAT1* promoter. Overall these results are consistent with a model in which HP1γ binding at promoter regions directly represses *STAT1* expression. Upon IFN-γ stimulation, HP1γ leaves the promoter to relieve gene expression.

## Discussion

Our work identifies a unique function for HP1γ in the control of the IFN-γ signaling cascade in the colon epithelium by repressing *STAT1* and *PD-L1* expressions. Depletion of HP1γ in the mice colon epithelium chronically de-repressed gene expression, pointing out a non-redundant and specific function of HP1γ with regards to others HP1 isoforms. To identify immune genes regulated by HP1γ, we overlapped the target genes for HP1 activity identified by RNA seq and the HP1γ ChIP seq data, thus revealing a group of genes under the control of IFN-γ signaling, including *STAT1* and *PD-L1*. Importantly, using intestinal organoid model and *in vitro* cell lines models, we showed that HP1 deficiency directly primes STAT1 and PDL1 expressions, providing seminal evidence for a silencing function on these genes at steady state level in the colon epithelium. ChIP assays further provided evidence that HP1γ directly repressed their expression, as shown by the constitutive HP1γ recruitment at genes promoters targetable upon IFN-γ treatment. Thus, our work supports a model in which HP1 exerts an epigenetic silencing at *STAT1* and *PD-L1*, whose detailed mechanisms deserve further investigations.

Remarkably, we found that HP1γ deficiency was sufficient to induce a chronic inflammatory state in the colon. This inflammation is characterized by T cell infiltrations and signs of disruption in epithelial homeostasis, including changes in goblet cells. These findings are reminiscent to those observed in the mucosa of UC patients and in mice models relevant for inflammatory bowel disease (IBD) (Tréton et al., 2014). Recent reports based on laser capture microdissection have provided evidence for high levels of IFN-γ in the lamina propria and surface epithelium of the colon mucosa in UC patients (Iacomino et al., 2020). Along with pro-inflammatory cytokines, anti-inflammatory cytokines such as IL-10 and TGF-β have also been detected in the inflamed intestinal mucosa of IBD patients (Autschbach et al., 1998). These findings suggest that immunoregulatory mechanisms are at play in IBD to reduce tissue damage and immunopathology. Interestingly, the IFN-γ signaling pathway has a dual nature, activating the innate immune response while also triggering immunosuppressive functions through the upregulation of PD-L1 and the differentiation of myeloid-derived suppressive cells (Lee and Ashkar, 2018). In this context, the control of PD-L1 expression by HP1γ may play a role in modulating intestinal tolerance in IBD and possibly in cancer immune escape (Han et al., 2020).

In conclusion, our work identifies HP1γ as an epigenetic silencing pathway controlling IFN-γ response at the epithelial barrier. We suggest that the previously identified decline in HP1γ in IBD patients (Mata-Garrido et al., 2022) might contribute to the deregulation of the immune responses by sensitizing the intestinal epithelium to IFN-γ signaling.

## Acknowledgment

HistIM Core Facility “Histology, Immunostaining, laser Microdissection”, Institut Cochin, Inserm U1016, CNRS UMR8104, Université Paris Cité.

## Materials and Methods

### Mouse model

The Villin-creERT2:*Cbx3*^-/-^ mouse model and Tamoxifen administration have been previously (Mata-Garrido et al., 2022)

### Colon crypts isolation and organoids culture

Colon crypt derived from 3 months aged male mice were isolated. Organoids were cultured in Matrigel as previously described (Mata-Garrido et al., 2022) After three days culture, 200U/ml mouse IFNγ (Miltenyi, 130-105-778) was added in the culture medium during 24h before RT-qPCR analysis.

### Cell culture and *CBX3KO* cell lines construction

SW480 and HT29 cells were cultured in Dulbecco’s modified Eagle medium (DMEM) supplemented with GlutaMAX (GIBCO, Life Technology) and 10% fetal bovine serum (FBS, Hyclone) with 5% CO2. To generate Crispr/Cas9-mediated *CBX3* KO cell line, HT29 and SW480 cells were transfected in 6-well plate in presence of Lipofectamine 2000 (Life Technologies), 1μg Cas9+sgRNA plasmid (Mata-Garrido et al. 2022) was used for each well. Single GFP^+^ cells were sorted by cytometry 48h after transfection. *CBX3* deletion in Crispr/Cas9 *CBX3*KO cell lines derived from different single cell clones were confirmed by Western blot and RT-qPCR before experiments.

### RT-qPCR

Total RNA was extracted using Trizol (TR-118, Molecular Research Center, Inc.) following the manufacturer’s instructions and DNAse treatment. RNA samples were quantified using a spectrophotometer (Nanodrop Technologies ND-1000). qPCR was performed using the Mx3005P system (Stratagene) with automation attachment. Primers used for samples are listed in **Table S1**.

### Western blot

Western blot performed from the CRC lysates were performed as previously described (Zhang et al. 2020). Mouse colon crypts were isolated and lysed in RIPA buffer (50 mM HEPES 0.1% SDS, 1% Triton X-100,1 mM EDTA,0.5% Sodium deoxycholate, 150 mM NaCl) with proteinase inhibitor (cOmplete™, EDTA-free Protease Inhibitor Cocktail, Roche). Western blot was realized with following antibodies: Anti-STAT1 (14994, cell signaling), Anti-Phosphorylated STAT1 (9167, cell signaling) Anti-HP1γ (IG-2MOD-1G6-AS, Euromedex), Anti-PD-L1 (4059, Prosci), Anti-Actin (A3854, Sigma), Anti-Tubulin (4D11, Thermo Scientific).

### Immunofluorescence and immunochemistry

Immunofluorescence was performed as previously described (Zhang et al. 2020) (Mata-Garrido et al. 2022). For organoids IF experiment, the organoids were cultivated on coverslip and fixed 4°C with 4% paraformaldehyde overnight before experiments. Anti-STAT1 (14994, cell signaling), Anti-ZO-1(Invitrogen, 33-9100) and Anti-HP1γ (IG-2MOD-1G6-AS, Euromedex) were used immune-labeling and DAPI was used for nuclear coloration.

For immunochemistry, slides were performed on the automaton Leica Bond RX and unmasked at pH 6 before incubated 30 minutes with an anti-CD8 antibody (Abcam, ab98941) or CD4 (Abcam, Ab183685), and then washed. The revelation system (“Bond Polymere Refine” kit, DS9800, Leica) included a secondary antibody HRP (Cell Signaling, 98941). Hematoxylin (blue) counterstaining allows the visualization of cell nuclei. CD4^+^ or CD8^+^ cells were counted with 4μM thickness colon tissue section with around 10 sections from 4 mice of each group.

### Flow cytometry analysis

Cells were detached with Trypsin-EDTA (Thermo Fisher, MA, USA), centrifuged and suspended in PBS containing 0.5% BSA, 2 mM EDTA and APC anti-mouse CD274 (Biolegend, 124312,) at 4°C for 30 min. After labelling, cells were washed once time and analyzed by LSR Fortessa™ cell analyzer (Becton Dickinson, NJ, USA).

### ChIP and ChIP-qPCR

ChIP experioments were performed as previously described (Saint-André et al., 2011). Clarified sample was incubated overnight with 2μg of HP1γ antibodies (Sigma-Aldrich, 05-690) or mouse non-immune IgG as negative control. Immunoprecipitated DNAs were quantified by RT-qPCR with primer sets specific for indicated CBX3 binding domain (Table S2, figure 6A and 6D). Values were measured relatively to the input and expressed relatively to the signal obtained for the immunoprecipitation with nonimmune IgG

### Bioinformatic pipelines

*CBX3* ChIP-seq dataset extracted from HCT116 (GSE192800) was mapped on human genome (HG38) with Binding and Expression Target Analysis (BETA) python algorithm and allowed identification of proximal binding sites around −3000 and +500 pb from the Transcription Stating Sites (TSS) of human genes (Wang et al., 2013). Gene set enrichment analysis extracted from CBX3 colon epithelium RNA-seq (GSE28115) was performed with GSEA version 4.3.0 standalone application (Han et al., 2020). During integrative analysis, HP1γ targets (ChIP-seq) found up regulated in RNA-seq were used as input for network analysis in STRING web tools version 11.5 (Szklarczyk et al., 2021).

### Statistical Analyses

Paired, unpaired or multiple two-sided t-test were performed with the GraphPad Prism based on at least three independent experiments or mice samples according to different experiment conditions. P-value inferior to 0.05 was considered as significant.

## Figures and legends

**Table S1:**
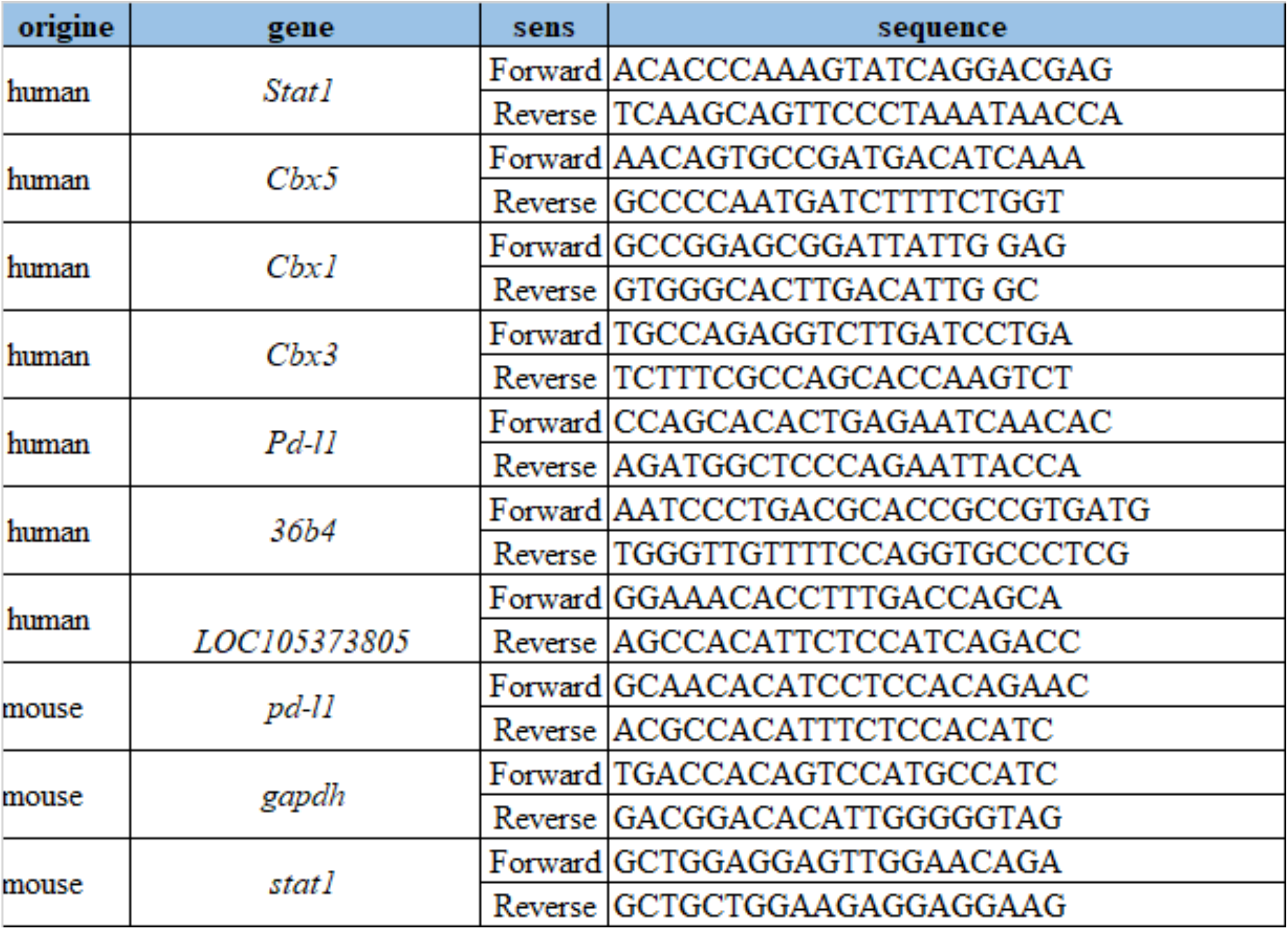
primer list for RT-qPCR

**Table S2:**
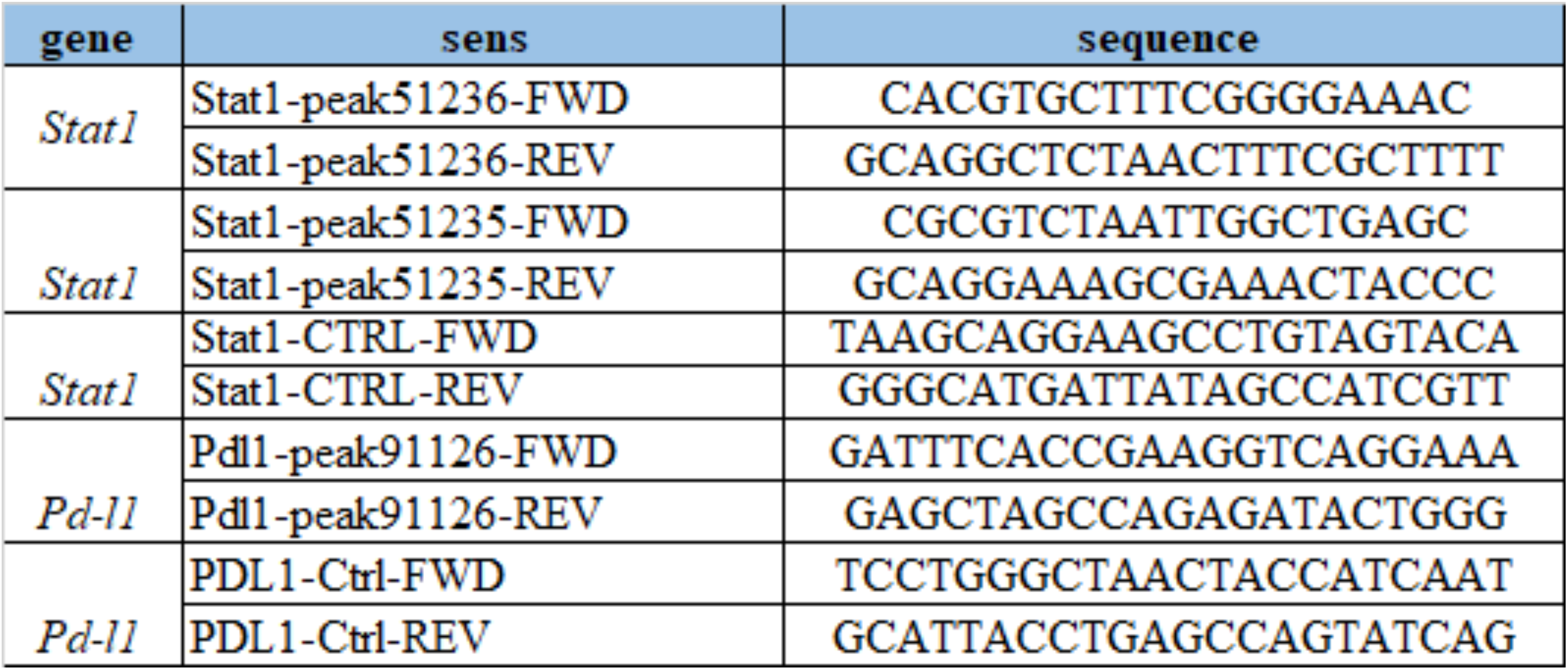
primer list for ChIP -qPCR

**Figure S1:**
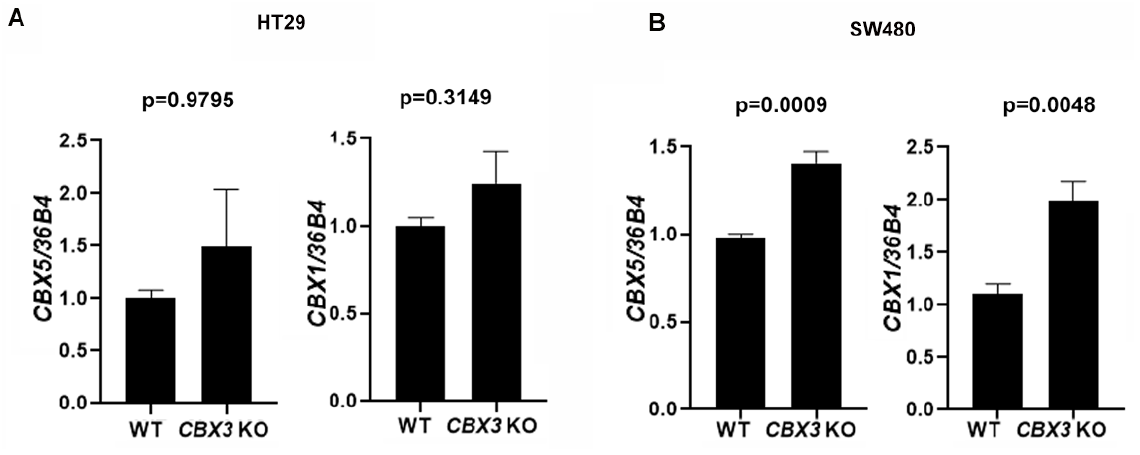
mRNA expression levels for *CBX5* and *CBX1* upon Cbx3 inactivation in the SW480 and HT29 CRC cell lines

**Figure S2:**
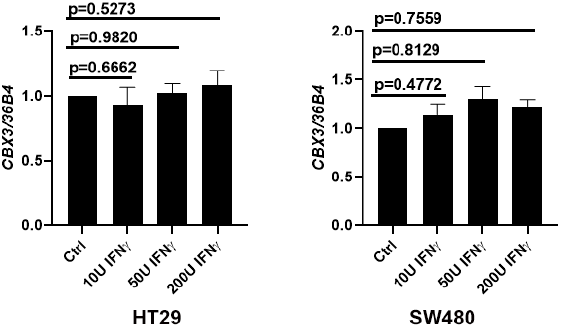
mRNA expression levels for HP1γ in response to IFNγ stimulations in the SW480 and HT29 CRC cell lines

